# Native Mass Spectrometry-Based Proteomics Reveals the Mechanism of Hemophore Release by Pathogenic *Corynebacterium diphtheriae*

**DOI:** 10.64898/2026.01.26.701814

**Authors:** Andrew K. Goring, Lindsey R. Lyman, Jordan Ford, Jingjing Huang, Christopher Mullen, Rafael D. Melani, Michael P. Schmitt, Robert T. Clubb, Joseph A. Loo

## Abstract

Iron is an essential micronutrient for nearly all forms of life, including pathogenic microbes that must acquire it from their host during infection. At the host–pathogen interface, humans restrict microbial access to iron through nutritional immunity, which many pathogens overcome by secreting hemophores that scavenge extracellular heme (iron protoporphyrin IX). However, identifying hemophores and other ligand-binding proteins in complex proteomes remains challenging using conventional peptide-based bottom-up mass spectrometry (MS). Here, we introduce ProteoMIX (Proteome Analysis by Mixing), a function-based native top-down proteomics workflow that combines slow-mixing mode native MS with charge reduction to identify ligand-binding proteins directly from complex mixtures. Applying ProteoMIX to the *Corynebacterium diphtheriae* exoproteome identified ChtA_30–314_, an abundant soluble hemophore generated by proteolytic processing of the surface-exposed ChtA heme receptor. Using native top-down MS sequencing, cell fractionation, and gene deletion, we show that the protease DIP2069 releases ChtA_30–314_ by removing ChtA’s transmembrane helix, producing a soluble proteoform that delivers heme to support microbial growth. In contrast, the related paralog ChtC is not processed, enabling *C. diphtheriae* to generate localization-specific heme-binding proteoforms from related gene products. Extending this approach to *Staphylococcus aureus*, ProteoMIX also revealed soluble IsdA hemophores that coexist with surface-anchored variants, demonstrating the generality of the method and suggesting that protease-mediated hemophore release may operate across gram-positive pathogens.

## Introduction

Iron is an essential micronutrient for virtually all forms of life, including pathogenic bacteria that must obtain it from their host during infection.^1^ At the host-pathogen interface, there is a battle for iron, with humans restricting microbial access to this metal through nutritional immunity mechanisms that pathogens attempt to overcome using a variety of iron-scavenging strategies.^2–4^ To capture iron in their surroundings, most pathogens secrete small-molecule siderophores that chelate iron and deliver it back to the cell to support growth.^5,6^ Pathogens also secrete hemophores that bind heme (iron-protoporphyrin IX) primarily derived from hemoglobin (Hb), which is a major reservoir for this nutrient as it contains ∼70% of the body’s total iron.^2,7–10^ While nearly all pathogens secrete siderophores, hemophores have been primarily discovered in gram-negative bacteria and appear to be rare in gram-positive pathogens.^5,7,11–19^ Despite their functional importance, identifying bacterial hemophores and elucidating the mechanisms through which they are released from the bacterium can be challenging. For example, *L. monocytogenes* Hbp2 is reported to be largely secreted with only a minor peptidoglycan-associated fraction, consistent with a hemophore-like role that complicates simple ‘cell-associated vs secreted’ classification.^20^ These challenges arise because bacterial genomes often encode several heme-binding proteins that, based on their sequences, are expected to remain cell associated.^21–25^ Moreover, their experimental detection can be cumbersome, requiring cell fractionation and the generation of appropriate antibodies to detect them by Western blotting.^26–29^ Bacteria can also secrete a range of other ligand-binding proteins, which can be challenging to detect using conventional methods (e.g., proteins that bind siderophores, metals, and other small molecules).^30,31^

*Corynebacterium diphtheriae* is the causative agent of diphtheria, a severe upper respiratory tract infection.^32^ To thrive in the iron-limited environment of its host, it produces a series of heme-binding proteins (HtaA, ChtA, ChtC, HtaB, and ChtB) that scavenge iron-laden heme and import it into the cell.^33–36^ Each protein binds heme via a Conserved Region (CR) domain and contains an N-terminal signal peptide for secretion through the Sec translocon and a C-terminal transmembrane (TM) helix that presumably embeds the proteins in the plasma membrane for display on the microbial surface. The microbe also displays a receptor (HbpA) that tethers Hb to the cell surface by contacting its α-globin.^22,37,38^ Surprisingly, peptide-based bottom-up proteomics and Western blot data indicate that the heme-binding ChtA and HtaA proteins are released from the cell into the exoproteome, despite the fact that their full length forms contain a TM helix for membrane association.^29,38^

More commonly employed MS methods are generally poorly suited for rapidly identifying specific ligand-binding proteins and their proteoforms within whole proteomes. For example, even though LC-MS/MS proteomics methods that measure proteolytically generated peptide fragments enable facile protein identification and quantification of their relative abundance, proteoform identification can be challenging compared to top-down proteomics.^39^ Moreover, native top-down MS (nTD-MS) methods can identify both proteoforms and their ligand-bound complexes, such as heme-bound hemophores. In this study, we present an nTD-MS approach that can rapidly identify ligand-binding proteins and their proteoforms within complex proteome mixtures without the need for extensive separation and fractionation. The method we call ProteoMIX (proteome analysis by mixing) obtains large gains in speed and efficiency by combining slow-mixing mode native MS (SLOMO-nMS)^40^ to monitor ligand binding with data-independent acquisition (DIA)-charge reduction to simplify the mass spectra.^41^ We demonstrate the utility of this method by discovering novel hemophores in pathogenic *C. diphtheriae* and *S. aureus* that are released by proteolytic processing of surface-associated heme-binding proteins. In *C. diphtheriae*, we define the mechanism of soluble hemophore release: the DIP2069 protease cleaves ChtA, removing its transmembrane helix and generating a soluble hemophore capable of scavenging heme from human Hb. Together, these results highlight ProteoMIX’s ability to identify ligand-binding proteins at the proteome level and uncover a previously unrecognized mechanism of hemophore release in Gram-positive bacteria.

## Results

### Native mass spectrometry enables the facile detection of hemophores within proteomes

Previously, we used bottom-up proteomics to identify 435 unique proteins within the *C. diphtheriae* exoproteome under iron-limiting conditions, which surprisingly included the heme-binding ChtA and HtaA proteins whose full-length forms contain membrane-interacting TM helices.^29^ This raised the possibility that ChtA and HtaA exist as truncated proteoforms in the exoproteome in which the TM helices were removed. In principle, native TD-MS provides a direct means to detect hemophores and other ligand-binding proteins; however, in complex proteomic samples the resulting spectra are highly congested with overlapping charge state envelopes that obscure proteoform assignments.^42^ To expedite the identification of functionally active hemophores and their proteoforms, we developed the ProteoMIX method and demonstrated its utility by applying it to the exoproteome of *C. diphtheriae*. ProteoMIX uses a two-step process to identify ligand-binding proteins, which in this application are hemophores that scavenge heme from human hemoglobin (Hb) (**Fig. 1**). In the first step, a plug of native exoproteome is loaded into an electrospray ionization (ESI) emitter tip followed by the addition of exoproteome that has been pre-incubated with Hb (**Fig. 1A**). This establishes slow-mixing mode native MS (SLOMO-nMS)^40^ between the exoproteome and analyte. A stable native electrospray is then established to generate a congested mass-to-charge (*m/z*) spectrum of the exoproteome whose intact masses are uninterpretable due to their convolved mass-to-charge ratios (**Fig. 1B**, left). Next, narrow *m/z* windows are isolated and charge-reduced by proton transfer- or electron capture-charge reduction (PTCR^43^ or ECCR,^44^ respectively) in a data-independent acquisition (DIA) scheme (**Fig. 1B**, middle). One cycle of DIA-charge reduction produces a snapshot of proteoform masses and relative intensities from the native ESI mass spectrum (**Fig. 1B**, right). Notably, DIA-PTCR on a Thermo Orbitrap Ascend Structural Biology Unit shows that 371 native proteoforms can be confidently detected from a single direct infusion (**Fig. S1**). This process is repeated as Hb diffuses from the back of the emitter tip to the front, and heme-binding proteoforms can easily be identified from intensity changes during the mixing period as they pick up heme (**Fig. 1C**). Proteoforms that bind heme lose intensity as their heme-bound proteoform, greater in mass by 616 Da (the mass of heme), gain intensity (**Fig. 1C**). Once hemophore masses are identified from the ProteoMIX dataset, they can be fragmented by nTD-MS (**Fig. 1D**). By using DIA-charge reduction coupled with SLOMO-nMS, proteomes containing hundreds of proteoforms (up to 371 in *C. diphtheriae*’s exoproteome in this example) can be screened for ligand-binding proteoforms in as little as one hour, and their most intense charge states can be prioritized for nTD-MS identification. This type of function-based prioritization of nTD-MS can reduce data collection time, as each proteoform identification requires non-trivial optimization in terms of data collection and analysis. Although we demonstrate the utility of ProteoMIX to identify heme-binding proteoforms, in principle it should be useful to study the binding of any ligand within a proteome that causes a measurable mass shift.

**Figure 1.**
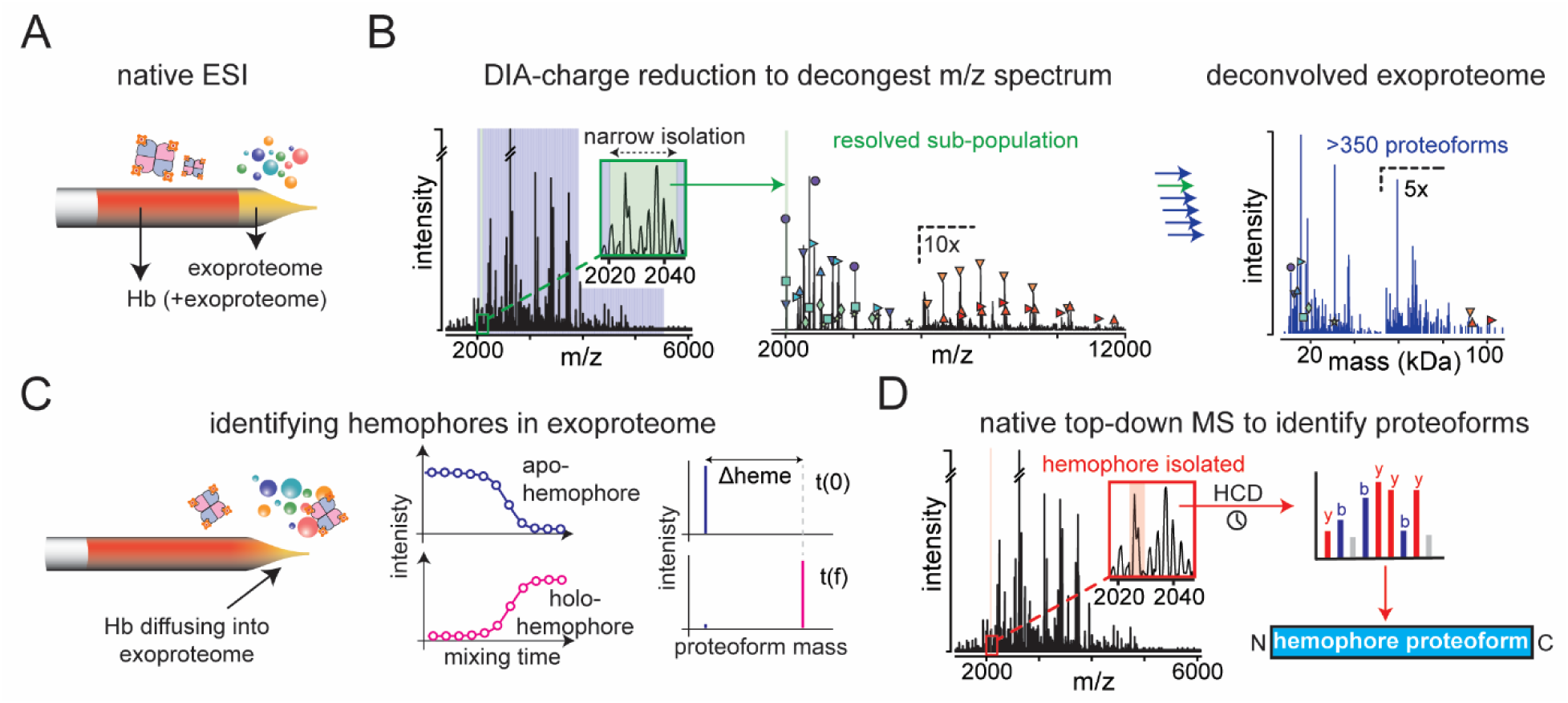
ProteoMIX-based approach for discovering ligand-binding proteoforms in unfractionated proteomes. (a) Buffer-exchanged exoproteome is loaded into the front of the nanospray tip followed by a stack of hemoglobin (Hb)-incubated exoproteome and a stable native ESI-MS spray is established. (b) Proteoforms in the resulting congested mass-to-charge spectrum are then resolved using DIA-charge reduction on narrow isolation windows spanning a majority of the exoproteome (e.g. 2000-5500 m/z, with 50 m/z spacing, for a total of 70 isolation windows). The charge-reduced spectra are deconvolved and summed in UniDec, and the resulting masses are evaluated by deconvolution score to produce a snapshot of proteoform masses and their intensities shown on the right in blue. (c) Hemophores are identified from changes in native proteoform intensities as hemoglobin slowly mixes into the exoproteome. Hemophore proteoforms are verified by paired mass shifts of 616 Da (Δheme). (d) Heme-binding proteoforms are then isolated based on their most abundant precursor mass-to-charge peak and fragmented using HCD to identify the hemophore’s proteoform.

### A truncated ChtA hemophore facilitates microbial growth by scavenging heme-iron

ProteoMIX reveals that *C. diphtheriae* releases from the membrane three fragments from the ChtA and HtaA proteins capable of scavenging heme: ChtA_30-314_, HtaA_31-272_, and HtaA_273-555_ (**Fig. 2A**). Each proteoform contains a heme-binding “conserved region” (CR) module, but lacks the C-terminal transmembrane (TM) helix that tethers the longer predicted secreted proteoform to the bacterium. An inspection of the N-termini in ChtA_30-314_ and HtaA_31-272_ reveals that they lack signal sequences, suggesting that the proteoforms originate from ChtA and HtaA proteins that have been secreted via the Sec translocon^45^. Heme capture by the highly abundant ChtA_30-314_ proteoform is readily observed from the 30,912 Da deconvolved mass gaining 616 Da (heme) during the mixing time-course (**Fig. 2B**). The full time-course tracking the intensities of apo- and holo-ChtA_30-314_ during mixing is shown in **Figure S2A**. nTD-MS reveals that the secreted ChtA_30-314_ hemophore contains an intramolecular disulfide bond between residues Cys179 and Cys188 (**Fig. S2B**), as previously observed in the crystal structure of ChtA’s CR domain (ChtA^CR^).^46^ HtaA_273-555_ was also identified from a clearly resolved 616 Da mass shift and reciprocal intensity changes for its apo-and holo-heme-bound proteoforms at 28,981 Da and 29,597 Da, respectively (**Fig. 2C**, **Fig. S2A**). Additionally, reciprocal intensity changes for the apo- and holo-HtaA_31-272_ proteoforms were crucial for its identification, as its apo-mass overlaps with the mass for secreted protease, DIP0350_30-274_, and therefore could have been missed without SLOMO-nMS (**Fig. 2D**). HtaA proteoforms were also verified by nTD-MS (**Fig. S2C**). Collectively, the data illustrate the utility of DIA charge-reduction combined with SLOMO-nMS for detecting ligand-binding proteins in complex mixtures and they reveal truncated ChtA and HtaA hemophores within the exoproteome of *C. diphtheriae* that are capable of scavenging heme.

**Figure 2.**
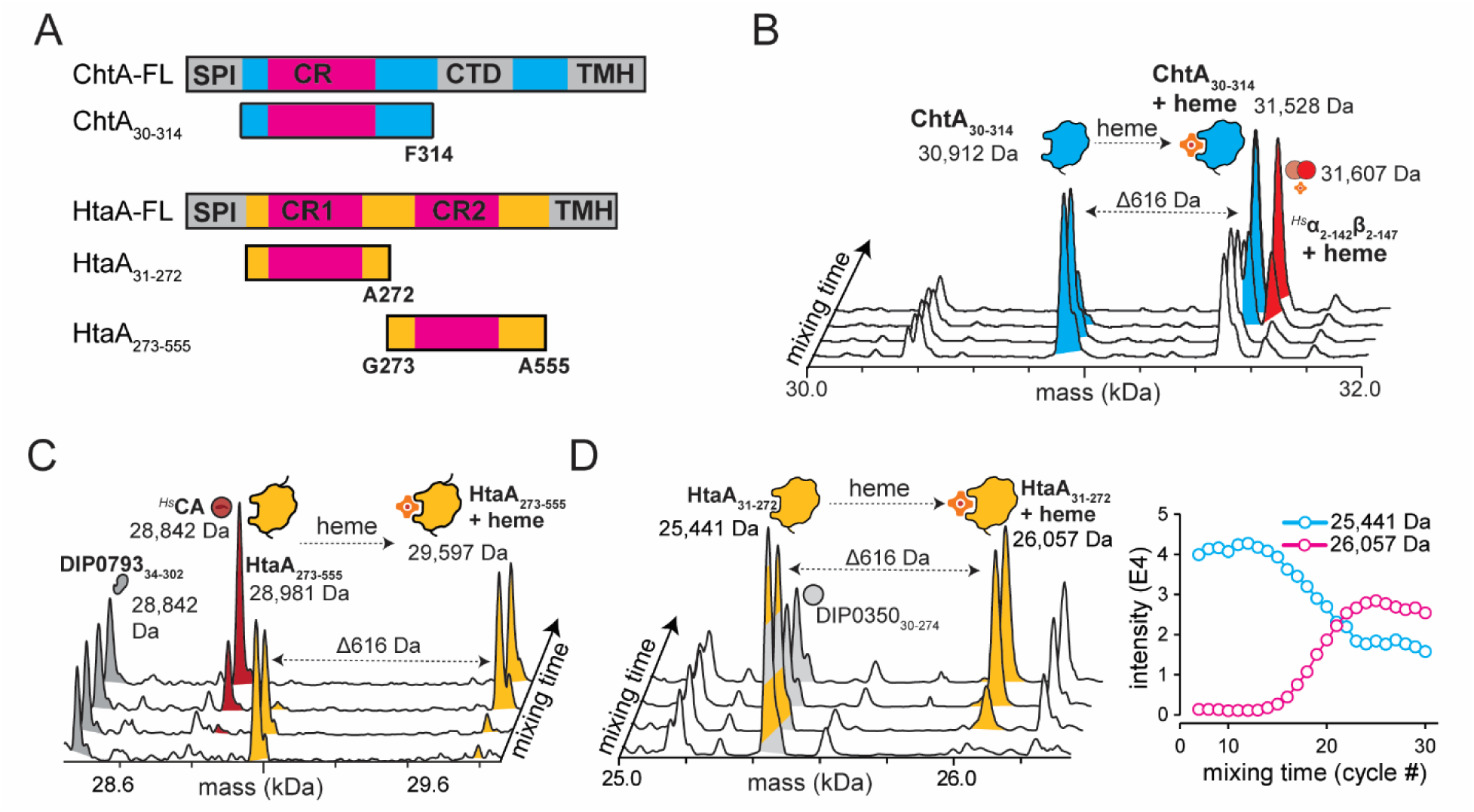
ProteoMIX identifies heme-binding proteins within *C. diphtheriae*’s exoproteome. (a) Full-length sequences with domains and sequence motif annotations for ChtA and HtaA (ChtA-FL and HtaA-FL, respectively). Hemophores identified using ProteoMIX (ChtA_30-314_, HtaA_31-272_, and HtaA_273-555_) are shown below the full-length sequences, and non-canonical termini are annotated below their illustration. (b) Select deconvolved mass spectra from DIA-charge reduction time points acquired using ProteoMIX show ChtA_30-314_ (blue) acquiring heme (Δ616 Da). The semi-apo Hb dimer (*^Hs^*α_2-142_β_2-147_ + heme) is also observed in red. (c) Analogous spectra show apo-HtaA_273-555_ binding heme (yellow; Δ616 Da). Trace amounts of human carbonic anhydrase (*^Hs^*CA) from the Hb sample are also observed diffusing into the exoproteome. (d) Deconvolved mass spectra (left) and extracted ion intensities from the full ProteoMIX time course (right) show the conversion of apo- to holo-HtaA_31-272_; reciprocal intensity changes were crucial for identification due to overlap with DIP0350_30-274_.

To determine the relative abundance of each ProteoMIX-detected hemophore, we performed bottom-up gel-enhanced LC-MS/MS proteomics (GeLC-MS). The exoproteome was separated by SDS-PAGE and then sectioned into 17 consecutive slices that were analyzed by LC-MS/MS (**Fig. 3A**). A total of 239 proteins were identified by GeLC-MS. Both full-length and truncated proteoforms of ChtA and HtaA are distinguishable and each exhibits a bimodal separation into low- and high-MW gel slices (**Fig. 3A**). When the molar abundance of all forms of each protein is summed, ChtA and HtaA are the 3^rd^ and 4^th^ most prevalent proteins in the exoproteome (at 8 ± 2% and 6 ± 1% molar abundance, respectively); the most abundant species is DIP2069, a peptidase S1 (clan PA) domain-containing protein (20 ± 8%), followed by the diphtheria toxin (10 ± 2%) (**Fig. 3B**). These data reveal that >99% of ChtA exists in its ChtA_30-314_ truncated proteoform, as nearly all of its intensity is located in the low-MW fraction. In contrast, peptides originating from HtaA are distributed between low- and high-MW fractions, revealing that only ∼30% of this protein exists as truncated HtaA_31-272_ and HtaA_273-555_ proteoforms, while the remainder is intact. Thus, secreted ChtA is predominantly present as the truncated ChtA_30-314_ proteoform. Given that ChtA_30-314_ is the most abundant heme-binding protein in *C. diphtheriae’s* exoproteome, we wondered whether its heme-bound form could serve as an alternate iron source to support bacterial growth. Indeed, exogenously supplied heme-loaded ChtA_30-314_ supports *C. diphtheriae* growth, as recombinant heme-loaded ChtA^CR^ domain restores growth in iron-limited media to levels comparable to cells using Hb as an iron source (**Fig. 3C**). Thus, the abundant truncated ChtA_30-314_ proteoform is not merely a degradation product, but a functional hemophore capable of supporting iron acquisition.

**Figure 3.**
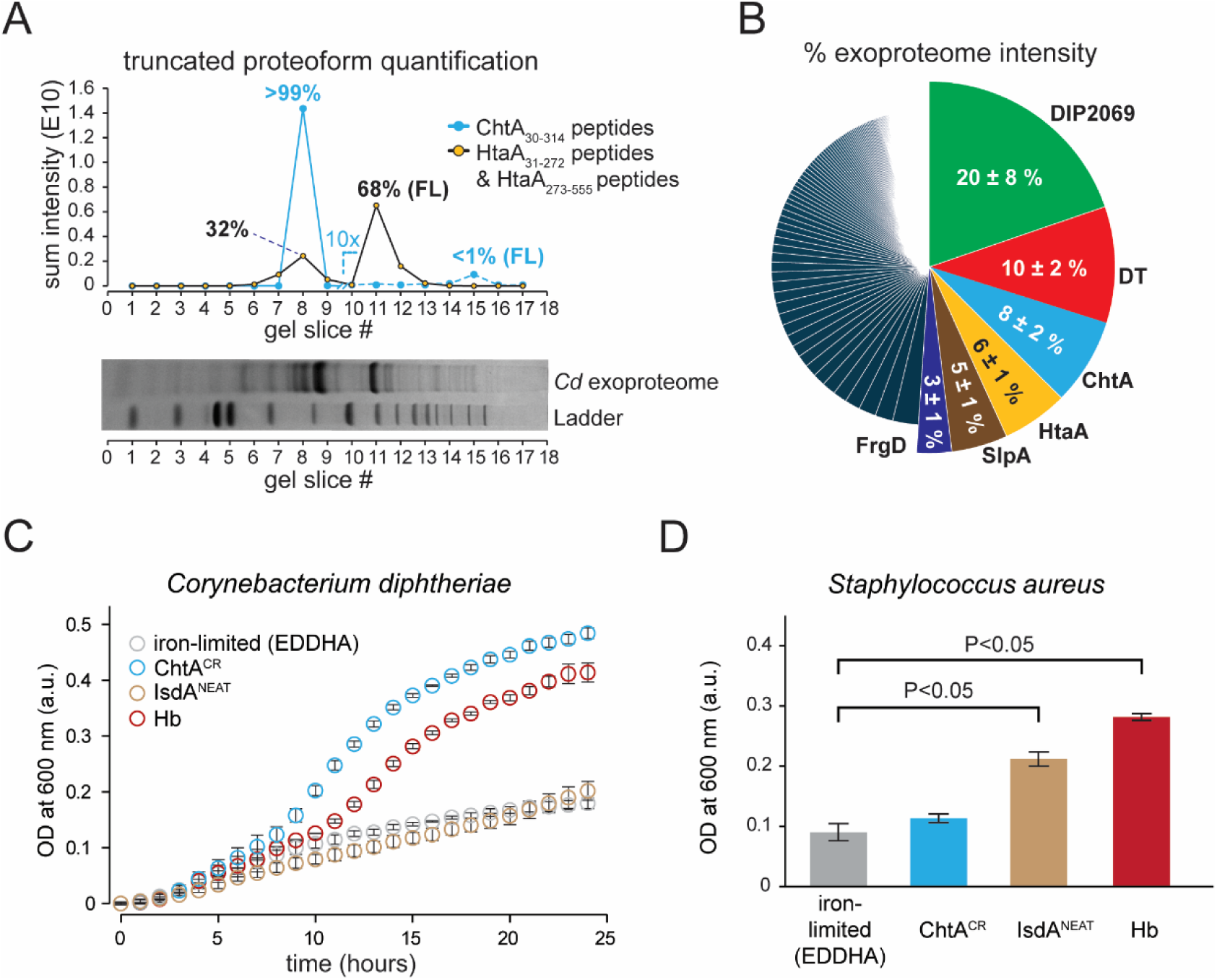
ChtA is a major exoproteome component, predominantly present as a truncated proteoform, and supports iron-limited *C. diphtheriae* growth through its heme-binding CR domain. (a) GeLC-MS on sequential SDS-PAGE slices shows ChtA is more than 99% present as a truncated proteoform, enriched in low-MW slices consistent with native MS. Only 32% of HtaA signal is present in low-MW slices. Intensities corresponding to full-length proteoforms are indicated with “FL”. (b) LC-MS/MS relative intensities of all proteins within the *C. diphtheriae* exoproteome show ChtA is highly abundant, following DIP2069 and DT. Relative intensities are in terms of iBAQ values (proxy for molar abundance) and correspond to full-length protein. Combining % intensity in (b) with % proteolysis from (a) provides an estimate of hemophore relative abundance. (c) Growth of *C. diphtheriae* on Hb and heme-loaded ChtA^CR^ at equivalent heme concentrations (0.6 µM) shows ChtA^CR^ supports growth better than hemoglobin. An evolutionarily distant heme-binding protein (heme-loaded IsdA^NEAT^ domain from *Staphylococcus aureus*) does not support growth to the same levels. (d) Growth of *S. aureus* in iron-poor media (RPMI 1640) supplemented with 250 µM iron chelator EDDHA and the same heme-loaded proteins at the same heme concentration used in (c) (0.6 µM) after 20 hours. Brackets indicate significant comparisons by two-tailed Student’s *t* test (P < 0.05).

### DIP2069 releases ChtA_30–314_ from the cell surface, whereas its ChtC paralog remains surface associated

Reasoning that ChtA and possibly HtaA are released from the cell surface by the action of a bacterial protease(s) that removes their C-terminal TM helices, we searched for other truncated proteins within the exoproteome that may also be processed by these enzymes. A prioritized target list of charge states in the raw spectrum was created for nTD-MS identification that corresponds to unique masses in the DIA-PTCR dataset. First, we collected 175 PTCR mass spectra from consecutively spaced isolation windows encompassing the bulk of ion current from direct infusion of the exoproteome (**Fig. S1**). Masses detected from the DIA-PTCR spectrum were then associated with their most abundant charge state in *m/z* space (**Fig. 4A**) and fragmented using higher-energy collisional dissociation (HCD) to reveal their primary sequence. Prioritization was critical for low abundance proteoforms. Without it, nTD-MS sequencing is prohibitively slow because each identification requires testing multiple fragmentation settings, each requiring long averaging times. The amino acid sequences from 60 proteoforms within the exoproteome were determined, which originate from 48 distinct proteins (**Table S1**, **Fig. 4A**). Interestingly, 7 of these 48 proteins correspond to novel truncated proteoforms resembling ChtA- and HtaA-like processing, as they contain C-terminal residues that are at either phenylalanine (ChtA, DIP2069, FrgD, RpfA) or alanine (HtaA, SprX, CtaC), respectively (**Fig. 4B**, **Table S1**). Specific cleavage sites within ChtA, HtaA and FrgD are positioned to remove C-terminal TM helix or lipid membrane anchoring elements, whereas cleavage of the other proteins presumably does not affect their location as their full-length forms are also expected to reside within the exoproteome. Compatible with their being processed by distinct proteases, the HtaA and DIP2069 proteins are cleaved at several sites that generate truncated fragments containing distinct C-terminal alanine and phenylalanine residues, respectively. Notably, all truncated DIP2069 proteoforms were observed with an additional ∼40 Da mass consistent with Ca²⁺ binding, supporting native cofactor engagement and a canonical trypsin-like serine protease fold for DIP2069 in the exoproteome (**Fig. S4**). Other proteoforms also appear in the exoproteome, and several of them show N-terminal cleavages corresponding to canonical signal peptidase cleavage sites (SpI and SpII) likely occurring after secretion through the Sec or Tat pathways (**Table S1**).^47^ Notably, ChtA_30-314_ is one of the most abundant truncated proteoforms within the exoproteome when assessed by native MS with DIA-charge reduction, whereas cleaved forms of HtaA (HtaA_31-272_, and HtaA_273-555_) are less prevalent because the full-length form of this protein predominates (**Fig. 3A**).

**Figure 4.**
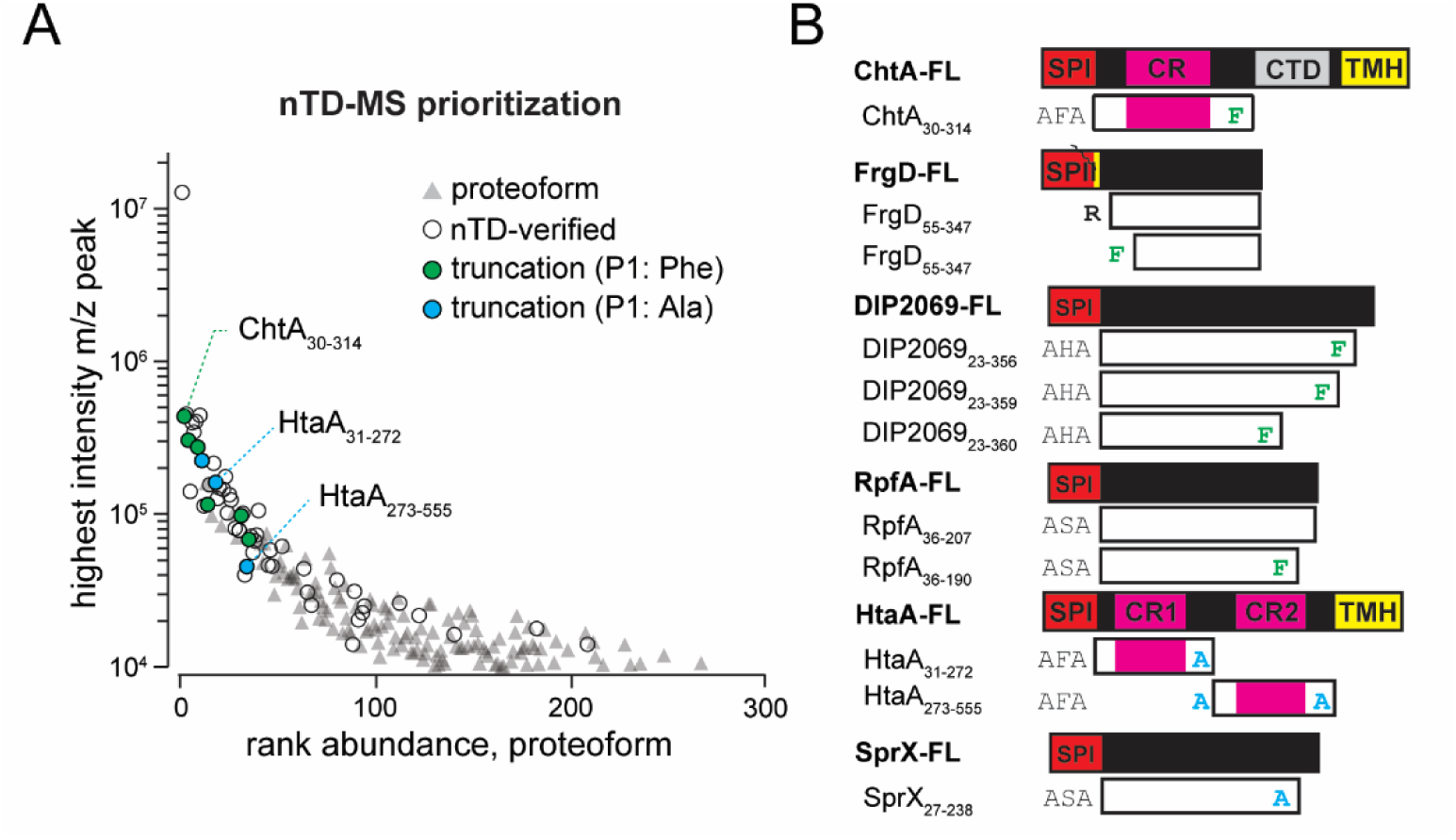
C-terminal phenylalanine cleavage reveals proteolysis of iron-acquisition proteins and exoenzymes. (a) Rank abundance of proteoforms detected by DIA-PTCR charge reduction, if they were identified by native top-down nMS (nTD-verified), heme binding (hemophore), and possess a non-canonical truncation site. The y-axis shows the intensity of each proteoform’s most abundant charge state in the DIA-PTCR dataset. (b) A total of 10 novel proteoforms from 6 different proteins of the exoproteome showed novel truncated proteoforms, 9 resulting from a C-terminal phenylalanine or alanine cleavage from their primary sequence. Two proteins involved in iron acquisition are missing membrane anchoring moieties which could explain their presence in the exoproteome (FrgD’s lipidated cysteine; ChtA’s C-terminal transmembrane helix). An exoprotease DIP2069 contains three novel C-terminal truncations at 3 closely spaced phenylalanine residues. RpfA which was found in its full length and truncated proteoform shows a C-terminal Phe cleavage.

To identify the protease responsible for releasing ChtA_30-314_ from the cell surface, we examined the effects of deleting the DIP2069 and SprX (DIP0964) extracellular proteases. They were chosen because they are abundant and exist as truncated proteoforms that contain either C-terminal phenylalanine (DIP2069_23-356_, DIP2069_23-359_, DIP2069_23-360_) or alanine residues (SprX_27-238_). The native exoproteomes of *C. diphtheriae* strains containing non-polar deletions in either *dip2069* (*Δdip2069*) or *sprX* (*ΔsprX*) were analyzed using DIA-charge reduction MS. While all three strains exhibit similar growth behavior in iron-replete media, the *Δdip2069* exoproteome differs markedly compared to the WT or *ΔsprX* deletion strain (**Fig. S3**). Notably, truncated ChtA_30-314_, among other truncated proteoforms is absent (**Fig. S3B**). Interestingly, Western blot analysis of the exoproteome shows that in the absence of *dip2069*, ChtA exists in its high-MW form in the exoproteome but this does not affect the high-MW form of HtaA (**Fig. 5A**, **Fig. S5**). We then performed label-free quantitative proteomics on exoproteome and surface fractions from WT and *Δdip2069* strains. Peptide intensities in the native exoproteomes of WT and Δdip2069 strains showed higher levels of peptides corresponding to the ChtA_30-314_ proteoform in the exoproteome, but significantly lower levels of the C-terminal region (**Fig. 5B**), further supporting that *dip2069* is responsible for forming ChtA_30-314_. Notably, in the absence of *dip2069*, ChtA is approximately 32-fold more abundant on the surface of the *Δdip2069* strain relative to WT (**Fig. 5C**). Thus, the native MS, western blot analysis, and LC-MS/MS proteomics data indicate that the DIP2069 protease liberates ChtA_30-314_ from the cell surface through proteolytic processing.

**Figure 5.**
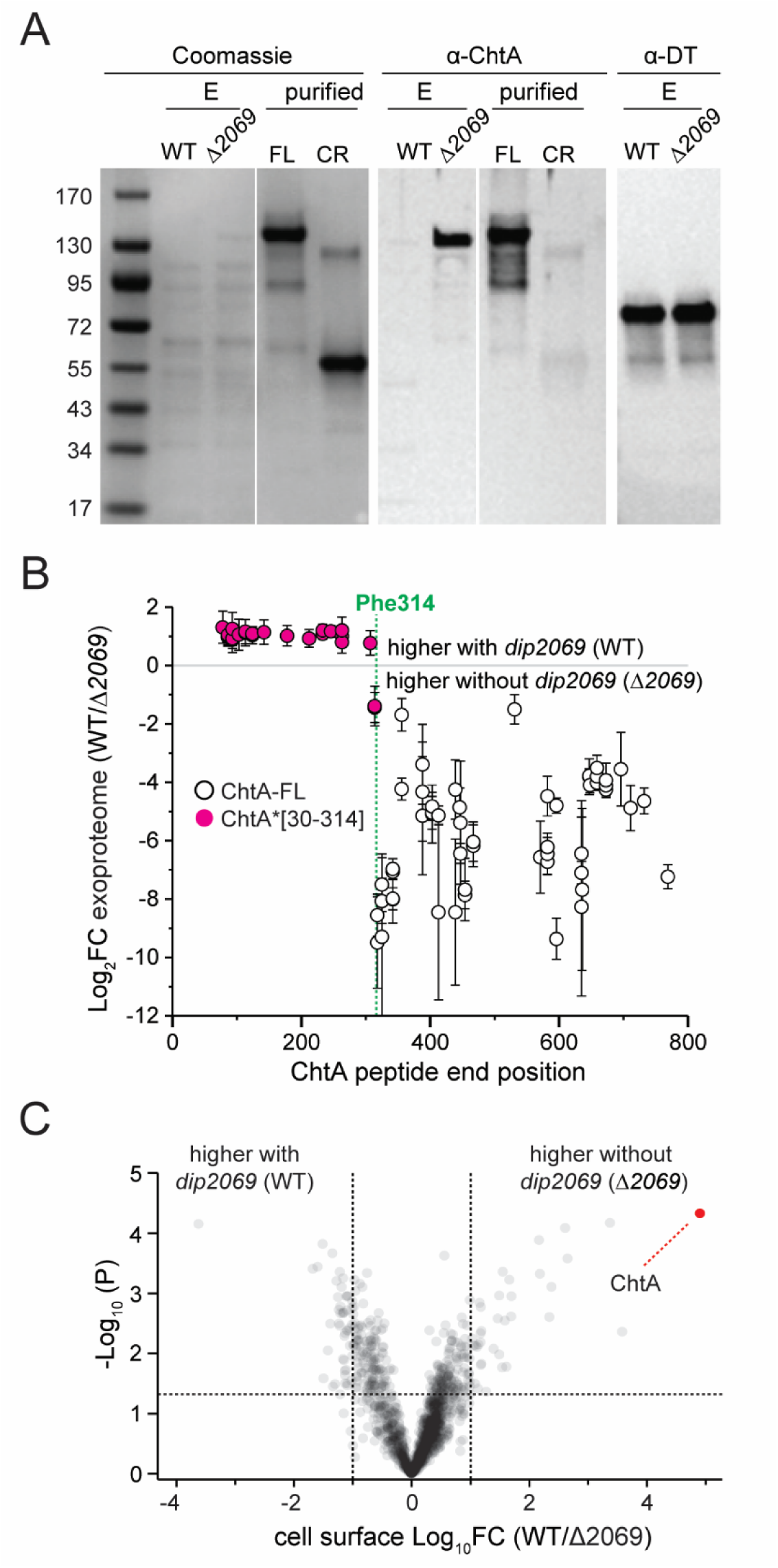
DIP2069 generates the secreted ChtA_30-314_ hemophore. (a) Cultures of iron-limited wild-type (“WT”) and *Δdip2069 C. diphtheriae* were grown to stationary phase and the resulting exoproteome (“E”) was subject to SDS-PAGE with Coomassie staining or prepared for western blotting with anti-ChtA (“α-ChtA”) or diphtheria toxin (“α-DT”) antibody. Additionally, recombinant full-length (FL) or ChtA encompassing the heme-binding CR domain with a C-terminal Strep-tag II (CR) was blotted with anti-ChtA antibodies to determine epitope coverage across ChtA’s sequence. (b) ChtA peptide intensity differences from exoproteome preparations of WT and *Δdip2069 C. diphtheriae* cultures. Only peptides with a significantly different abundance between each sample as defined by a P-value of 0.05 from an unpaired student’s T-test are shown. (c) A volcano plot showing abundance differences at the cell surface between WT and *Δdip2069* strains from the same cell cultures that exoproteomes were prepared from for the data in (b).

In addition to ChtA, *C. diphtheriae* also expresses the closely related ChtC protein which shares 47% sequence similarity.^48^ Both contain related heme-binding CR domains and C-terminal helices, however, ChtA and ChtC are unique members of the heme uptake machinery because they are also predicted to harbor a central beta-propeller domain whose function is not known. Interestingly, the MS data reveal that these proteins localize to distinct sites – ChtA partitions to the exoproteome as ChtA_30-314_, whereas ChtC remains associated with the microbe and exposed on the cell surface (**Fig. 6B**).^29^ An inspection of their primary sequences suggests that this difference may be a result of distinct susceptibilities to DIP2069 cleavage, as the phenylalanine residue that is the cleavage-site in ChtA is conserved across species, whereas in the *C. diphtheriae* ChtC protein this residue is replaced with a tryptophan (**Fig. 6A**). Indeed, compatible with it not being a substrate for DIP2069 in both WT and *Δdip2069* strains the ChtC protein remains cell-associated (**Fig. 6C**), whereas *Δdip2069* deletion prevents formation of ChtA_30-314_ (**Fig. S3B**) in the exoproteome and leads to accumulation of ChtA on the cell surface (**Fig. 5C**). Together, the data suggest that DIP2069 is responsible for proteolytically releasing ChtA_30-314_ from the cell surface, whereas a subtle sequence alteration in its paralog ChtC causes it to remain associated with the cell surface.

**Figure 6.**
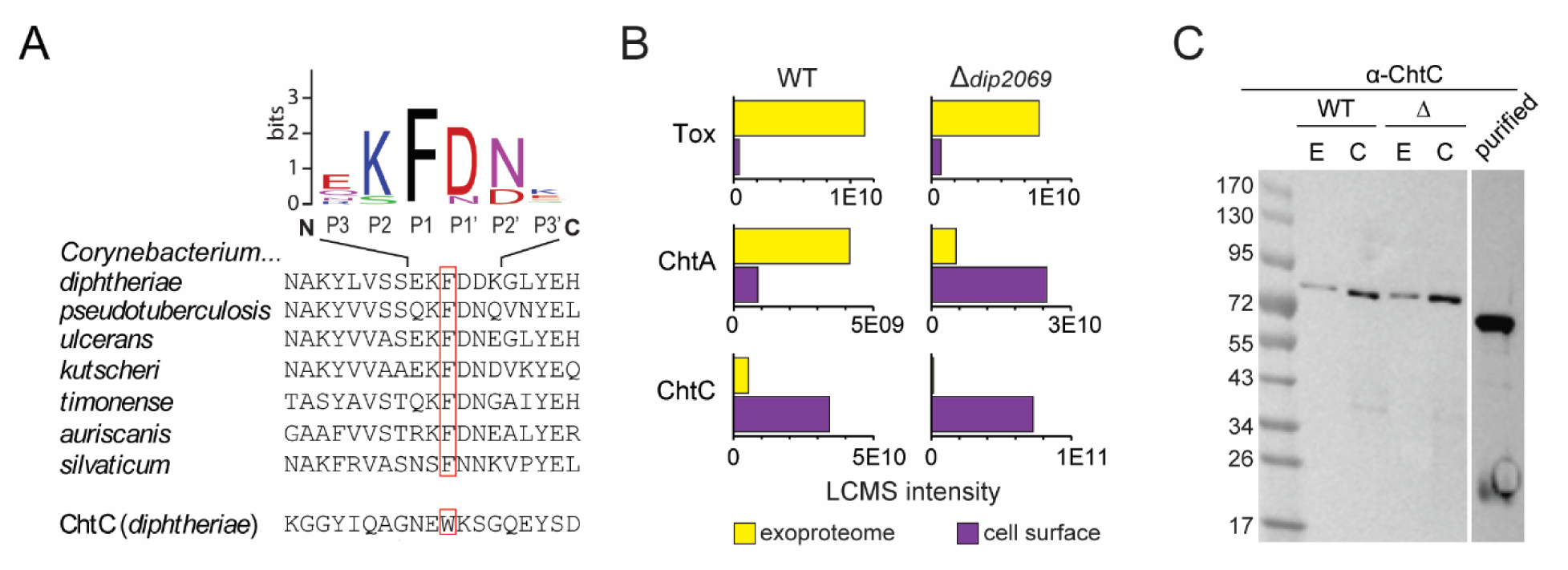
DIP2069 selectively liberates the ChtA_30-314_ hemophore from the cell surface, but not its paralog ChtC. (a) Sequence alignment of ChtA-like proteins across prominent corynebacterium pathogens show Phe314 is highly conserved and flanked by charged and/or polar residues. Similarly, polar and charged residues are present in DIP2069, FrgD, and RpfA novel proteoforms that arise from cleavage events C-terminal to phenylalanine. (b) Peptide intensity differences from exoproteome and cell surface preparations of WT and *Δdip2069 C. diphtheriae* cultures. (c) Western blot analysis of ChtC in the exoproteome (“E”) and cell surface (“C”) fractions from WT and *Δdip2069* strains showing full-length ChtA across all fractions.

### ProteoMIX reveals *S. aureus* releases IsdA for heme scavenging

To demonstrate its broad utility, we used ProteoMIX to characterize the exoproteome of pathogenic *S. aureus*. Previous studies have shown that this bacterium expresses heme-binding Iron-regulated Surface Determinant (Isd) proteins that are covalently attached to the peptidoglycan (IsdA, IsdB, IsdC, and IsdH).^49^ Application of ProteoMIX revealed soluble IsdA proteoforms in the exoproteome capable of scavenging heme released from Hb (**Fig. S6A, B**). Similar to *C. diphtheriae* ChtA_30-314_, these IsdA proteoforms (IsdA_46-224_, IsdA_46-226_, and IsdA_46-238_) lack C-terminal segments that tether the full-length, cell wall-associated protein to the microbe. Three truncated IsdA proteoforms are generated by cleavage of the segment connecting the heme-binding Near Transporter (NEAT) domain to the C-terminal LPXTG motif (**Fig. S6A**), which covalently attaches the surface-associated protein to the peptidoglycan.^50^ GeLC-MS studies indicate that IsdA partitions roughly equally between cell wall-associated and extracellular forms.^51^

*S. aureus* and *C. diphtheriae* can sometimes inhabit overlapping sites in the upper respiratory tract, and the two bacteria are common co-infectants in subcutaneous wounds,^52^ suggesting their hemophores may compete for heme iron to sustain growth. Interestingly, growth assays show that heme delivery to *C. diphtheriae* is specific for ChtA, as supplementation of iron-limited *C. diphtheriae* cultures with the recombinant heme-loaded form of the structurally distinct IsdA^NEAT^ hemophore from *S. aureus* does not appreciably rescue growth (**Fig. 3C**). A similar result is observed for *S. aureus* under iron-deplete conditions. The bacteria grow when supplemented with either Hb or recombinant heme-loaded IsdA^NEAT^, whereas heme-loaded ChtA^CR^ from *C. diphtheriae* is largely ineffective in supporting *S. aureus* growth in iron-limited media (**Fig. 3D**). These results indicate that productive heme delivery is largely species-specific, with each hemophore preferentially supporting growth of its native bacterium.

## Discussion

Protein-ligand binding events are fundamental to nearly every aspect of cellular function (catalysis, signaling, growth, and metabolism among others), yet identifying these interactions within complex proteomes remains challenging.^53^ Here we report the development of ProteoMIX, a nMS approach that efficiently identifies specific ligand-binding proteins and their proteoforms within proteomic mixtures. ProteoMIX builds on two recent advancements in native MS: (i) slow mixing mode (SLOMO) nMS that quantitatively tracks gradual changes in protein intensities as a ligand binds,^40,54,55^ and (ii) DIA-charge reduction that dramatically reduces spectral complexity (**Fig. 1**).^41,43,44^ By integrating these methods, we identified heme-binding proteoforms directly from the unfractionated exoproteomes of *C. diphtheriae* and *S. aureus*, shedding new light on how these pathogens scavenge heme during infections. SLOMO was used to track heme uptake from Hb by this microbe’s exoproteome with simultaneous DIA-charge reduction to deconvolve the mass spectra. We show that heme-binding proteoforms can be identified with high confidence by paired losses and gains matching the mass of heme, enabling the prioritization and ensuing top-down MS sequencing of their most abundant charge states (**Fig. 2**). The prioritization step greatly improves efficiency for hemophore identification, as the alternative would be to fragment every peak in the native mass spectrum. Given current throughput, confident MS2 identification of low-to-moderate abundance proteoforms is slow, making comprehensive peak-by-peak sequencing impractical currently (often hours).

Using ProteoMIX we discovered that *C. diphtheriae* secretes ChtA_30-314_ to scavenge heme from Hb and other hemoproteins. This was surprising, since like other proteins in *C. diphtheriae*’s heme uptake system that are instead cell wall associated, the full-length ChtA protein contains a C-terminal TM helix that is presumably embedded in the bilayer to enable surface display. In follow-up experiments that used a combination of GeLC-MS (**Fig. 3**), nMS (**Fig. S3B**), and Western blotting applied to wild-type and *Δdip2069* strains (**Fig. 5**, **Fig. 6**), we provide evidence that a highly abundant extracellular chymotrypsin-like protease DIP2069 is responsible for releasing ChtA_30-314_ from the cell. ChtA_30-314_ is liberated when the surface-displayed full-length ChtA protein is cleaved by DIP2069 between its heme-binding CR domain and its C-terminal propeller domain. Once outside the cell, ChtA_30-314_ can capture heme and deliver it back to the bacterium to enable growth when iron is scarce (**Fig. 3C**), suggesting it plays an important role in iron uptake during infection. Other pathogenic corynebacterial bacterial species presumably use a similar protease-mediated release mechanism to release soluble ChtA hemophores for iron scavenging, as they also contain DIP2069 homologs and ChtA proteins that share a conserved sequence surrounding the site of cleavage. A systematic analysis of the nMS spectra reveals several other truncated proteoforms in the exoproteome that may be generated by DIP2069, as they share conserved C-termini with ChtA_30-314_ (FrgD, an iron-siderophore/vitamin B12 binding protein; RpfA, a growth resuscitation factor; and DIP2069 itself) (**Fig. 4B**).

By leveraging unbiased proteomic profiling, we identified HtaA as a second secreted hemophore, present in both full-length (**Fig. S5**) and proteolytically cleaved proteoforms (**Fig. 2**, **Fig. S2**). ProteoMIX detected two HtaA proteoforms (HtaA_31-272_ and HtaA_273-555_) in the exoproteome, which each contain one of its two CR domains (CR1 and CR2), but lack the TM helix present in the parent protein. These proteoforms are presumably generated by a yet-to-be-identified protease, as HtaA abundance does not change substantially in the *Δdip2069* strain (**Fig. S5**), and the C-termini of HtaA_31-272_ and HtaA_273-555_ lack the C-terminal phenylalanine residue likely present at the DIP2069 cleavage site. Notably, in contrast to ChtA, which is almost completely converted into ChtA_30-314_ by the DIP2069 protease, only ∼30% of HtaA is cleaved. Outside the cell, the longer TM-helix-containing HtaA proteoform predominates, whereas HtaA_31-272_ and HtaA_273-555_ are roughly five times less abundant than ChtA_30-314_. Interestingly, the longer HtaA proteoform is undetectable by nMS, consistent with it being insoluble or sequestered in mycomembrane vesicles too large for detection. This idea aligns with recent findings that the related bacterium *Corynebacterium glutamicum* produces mycomembrane vesicles containing its heme-binding proteins.^56^ Interestingly, large HbpA aggregates have previously been observed in the exoproteome,^37,57^ raising the possibility that it may also be packed into vesicles.

Our findings clarify the roles of individual components within the *C. diphtheriae* heme-uptake system and reveal distinct functional specializations of its paralogous ChtA and ChtC proteins. **Figure 7** summarizes the positioning of these protein components based on our results and previously reported protease shaving and cell fractionation studies.^58^ HbpA and ChtC are exposed on the mycomembrane, where they are positioned to bind Hb and heme, respectively. Also exposed on the surface are portions of the heme-binding ChtA and HtaA proteins, which are significantly less abundant than HbpA and ChtC; the approximate abundance ratios are 10:5:1:1 for HbpA:ChtC:ChtA:HtaA. Each of these exposed proteins is presumably tethered to the underlying plasma membrane via their C-terminal TM helices, with N-terminal ligand-binding domains positioned outward. HtaA is unique in containing two heme CR domains, CR1 and CR2. Considering the dimensions of the corynebacterial envelope, CR1 is likely exposed on the surface, whereas the C-terminal CR2 domain is buried within the peptidoglycan. The heme-binding HtaB and ChtB proteins are also buried within the peptidoglycan and tethered to the membrane via C-terminal TM helices. Outside the bacterium, our results show that ChtA_30-314_ scavenges heme from Hb to facilitate cell growth after it is proteolytically released from the cell surface by extensive cleavage of surface-embedded ChtA by the DIP2069 protease (the majority of ChtA is cleaved, such that ChtA_30-314_ is ∼32 times more abundant than cell-surface ChtA) (**Fig. 5**). Interestingly, even though ChtC and ChtA share highly similar primary sequences,^48^ ChtC is a poor substrate for DIP2069 and remains intact on the cell surface (**Fig. 6**). At least three heme-scavenging proteoforms of HtaA are also present extracellularly: a nearly intact proteoform containing its TM helix and both heme-binding domains (**Fig. 7**), and two less abundant proteolytic fragments, each containing a single heme-binding domain (HtaA_31-272_ and HtaA_273-555_) (**Fig. 2**). Thus, in its quest for iron, *C. diphtheriae* releases and displays an array of proteins to scavenge heme, presumably delivering it to the microbial surface for transfer across the cell envelope. How heme is transferred among these components is not known, but transfer studies using a fluorogenic reporter suggest it may occur via transient heme-transfer complexes involving their CR domains.^59^

**Figure 7.**
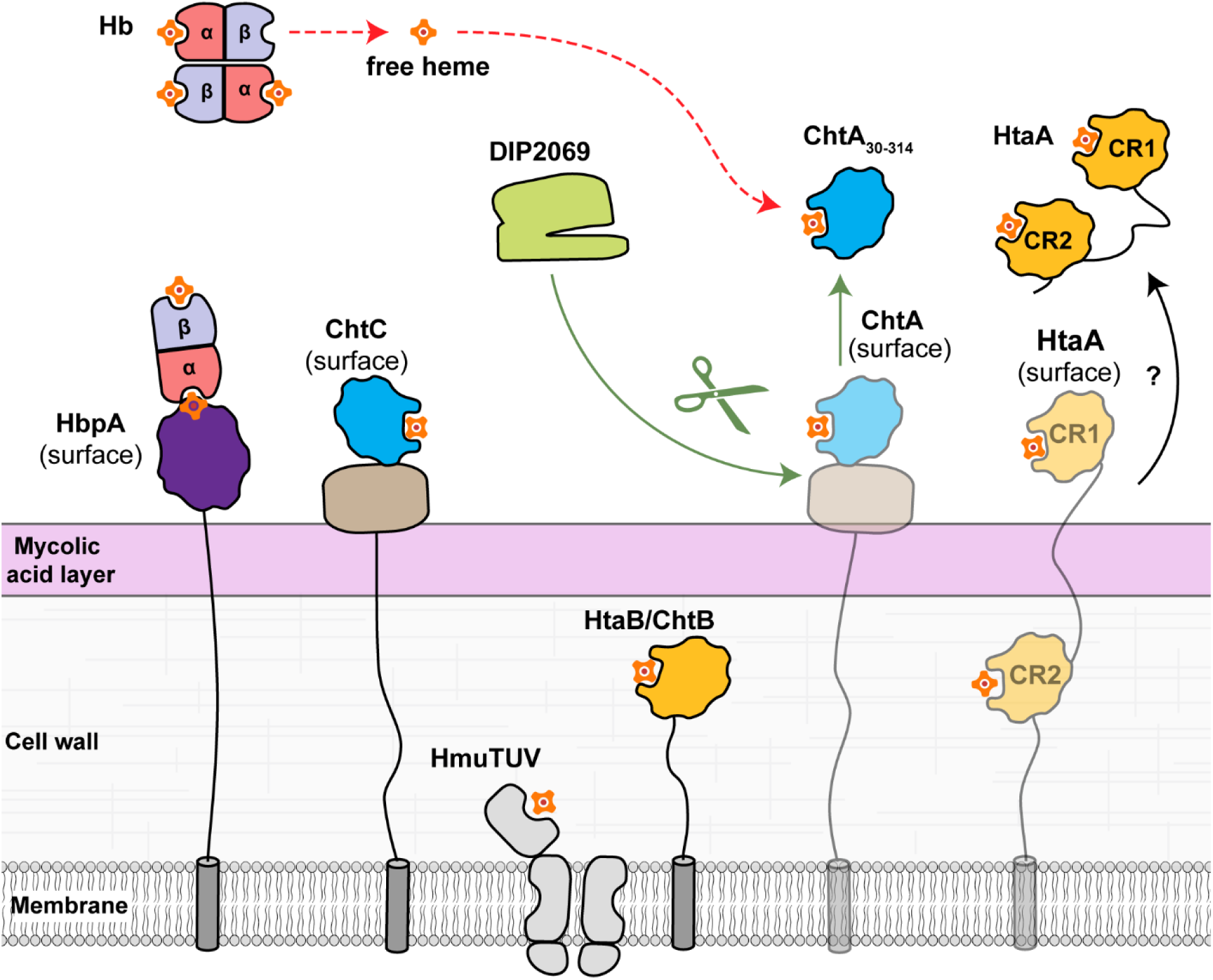
A model of protease-mediated heme acquisition by *Corynebacterium diphtheriae*. Heme-acquisition begins with red blood cell lysis near the site of infection, releasing high concentrations of tetrameric hemoglobin. Tetrameric hemoglobin releases free heme and dissociates into dimeric hemoglobin, which release heme approximately 8 times faster. The free heme is scavenged by host hemopexin and *Corynebacterium diphtheriae*’s secreted ChtA_30-314_ hemophore, which it ultimately uses as an iron source. The ChtA_30-314_ hemophore is produced from surface-associated full-length ChtA, which DIP2069 selectively processes, in contrast to ChtA’s surface-associated paralogue, ChtC, which remains on the cell surface during iron-limited conditions. Heme bound to ChtA_30-314_ is the passed to *C. diphtheriae* to obtain its iron. This presumably occurs when it transfers heme to cell associated HtaA, HtaB and ChtB heme-binding proteins, or directly to the HmuT for import by the HmuTUV ABC transport cassette.

Application of ProteoMIX to evolutionarily-distinct gram-positive *S. aureus* revealed that this pathogen also releases soluble hemophores (**Fig. S6**). We identified truncated IsdA proteoforms in the exoproteome that coexist with longer, cell wall-anchored variants, revealing a modular system in which surface-bound IsdA proteins capture local heme while secreted proteoforms presumably extend acquisition range. The parallel processing of IsdA in *S. aureus* and ChtA_30-314_ in *C. diphtheriae* suggests that proteolytic liberation of hemophores is a conserved strategy among Gram-positive bacteria. Growth assays highlight species-specific functionality: while heme-loaded ChtA_30-314_ delivers heme to *C. diphtheriae*, delivery by the structurally unrelated IsdA^NEAT^ hemophore occurs far less efficiently. The converse also is true; exogenously supplemented heme-loaded IsdA^NEAT^ is capable of delivering heme and sustaining microbial growth of *S. aureus* for better than ChtA^CR^ (**Fig. 3D**). This suggests that these hemophores have coevolved with their cognate uptake machinery to provide a competitive growth advantage in nutrient-limited niches colonized by diverse genera of bacteria. Notably, *C. diphtheriae* reaches a higher optical density when supplied with heme pre-loaded onto ChtA^CR^ than with an equivalent concentration of Hb, consistent with hemophore-mediated delivery bypassing passive Hb-heme extraction on the cell surface. In contrast, *S. aureus* grows slightly better on Hb than on IsdA^NEAT^, reflecting the presence of dedicated Hb receptors (e.g., IsdB and IsdH) that efficiently harvest heme from Hb.^60–62^ In contrast, *S. aureus* grows slightly better on Hb than on IsdA^NEAT^, aligning with the presence of dedicated Hb receptors (e.g., IsdB/IsdH) that can efficiently harvest heme from Hb in an active process.^60–62^ These observations suggest that secreted IsdA proteoforms in *S. aureus* may primarily extend the range of heme scavenging or sequester heme to limit access by competitors, rather than serving as the main mechanism for heme acquisition from Hb. Because relatively few Gram-positive hemophores have been identified to date, many more may be uncovered using ProteoMIX. Collectively, these findings establish ProteoMIX as a powerful platform for discovering functional hemophores, revealing previously hidden layers of bacterial nutrient acquisition, and opening new avenues for targeting iron metabolism in pathogens.

## Acknowledgments

The authors would like to thank members of the Clubb and Loo labs for useful discussions. This work was supported by grants from the NIH (R35GM145286 to J.A.L, R01AI161828 to R.T.C.). A.K.G. acknowledges support from the NIH NIDCR (T90DE030860). Additional support was provided by the NIH National Institute of General Medical Sciences (NIGMS)-funded predoctoral fellowship for J.F. (T32GM145388).

