## Supplementary Information for "Native Mass Spectrometry-Based Proteomics Reveals the Mechanism of Hemophore Release by Pathogenic *Corynebacterium diphtheriae*"

**Materials and Methods** ........ **S2**

**Supplementary Figures** ........ **S6**

**Supplementary Tables** ........ **S12**

**References** ........ **S14**

#### Materials and Methods

**Strain and materials**. *Corynebacterium diphtheriae* strain 1737 was used in this work. Non-polar deletions of *dip2069*, *dip0775*, and *sprX* (*dip0964*) were also made from this strain. For all cell cultures, glycerol stocks were used to form single colonies on an heart infusion agar plate were used to inoculate starter cultures of *C. diphtheriae* in 5 mL HIBTW (heart infusion broth + 0.2% tween-80).

**Exoproteome preparation for native MS**. Exoproteome was made from cells cultured in mPGT media without iron supplementation. Briefly, starter cultures were diluted 1:2 in HIBTW (HIB + 0.2% tween-80) and cultured for 1 hour. The resulting culture was centrifuged and washed once in fresh mPGT media. The cultures were then inoculated into mPGT media, 3x 15 mL cultures. Each 15 mL culture was incubated without shaking, in an agar plate to increase surface-to-volume ratio. These cultures were filtered, incubated with protease inhibitor cocktail (Peirce), and concentrated to 100 µL using a 10 kDa cut-off filter. This was then subject to 2 rounds desalting into 200 mM ammonium acetate using Zeba 7 kDa MWCO 0.5 mL desalting columns (Fischer) and desalted until optimal signal intensity using an Amicon 10 kDa MWCO centrifugal filter. Incubations of this with or without hemoglobin in stacked layer format

**Native MS**. Native exoproteome was sprayed on either a Thermo Scientific Q-Exactive series Ultra High Mass Range (UHMR) mass spectrometer or a Thermo Scientific Orbitrap Ascend BioPharma Tribrid mass spectrometer with Native MS option. Native electrospray was conducted with borosilicate nanospray capillaries coated with platinum, at a spray voltage of 0.96 kV. Charge reduction was performed with electron capture charge reduction (ECCR) when using a UHMR, and proton transfer charge reduction (PTCR) when collected on an Orbitrap Ascend.

**Cell fractionation for bottom-up proteomics and western blotting**. Iron-limited early stationary-phase culture supernatant and cell pellets from wt- and DIP2069-devoid cells were used as the starting material for cell surface vs secretion localization profiles from bottom-up proteomics. The supernatants were prepared for bottom-up proteomics following concentration, reduction, alkylation and trypsinization. Cell surface fractions were prepared by incubating the cell pellets from the resulting cultures with LCAO and beta-octyl glucoside, shaking for 37 C for 1 hr, pelleting cell debris, and removing the supernatant.

**Western blotting**. Cells used for gel electrophoresis and Western Blotting were grown in mPGT medium, prepared as described previously[1] with $FeCl_3$ supplementation at 0.25μM and normalized by OD600 at harvest. After reaching stationary phase, cultures were centrifuged and supernatant harvested for exoproteome assessment. Normalized quantities of supernatant were boiled for 10 minutes in Laemmli buffer and separated on 4-15% denaturing precast TGX (Tris-Glycine eXtended) PAGE gels using tris-glycine-SDS running buffer (all reagents from Bio-Rad). Coomassie staining, transfer, and western blot procedures were done as previously described.[2] Anti-ChtA and anti-DT primary antibodies were used at 1:10,000.

Purified proteins for Western Blots were prepared as follows: Recombinant strep-tag II-tagged proteins were expressed from a pET24(a)+ vector and purified from *E. coli* strain BL21(DE3) following growth in Luria-Burtani medium and induction at 27°C for 3 hr with Isopropyl β-D-1-thiogalactopyranoside (IPTG). For ChtA, the construct comprised the ChtA CR domain (ChtA residues 110–299) fused to a C-terminal Strep-tag II. Cells were lysed using the FastPrep cell lysis system (MPBiomedical) followed by centrifugation for 10 min at 4°C. Lysis was done in buffer W (100 mM Tris-HCl [pH 8.0], 150 mM NaCl) as recommended by the manufacturer (IBA Lifesciences) for purification of strep-tagged proteins. Streptactin Sepharose columns (IBA Lifesciences) were used with the manufacturer’s instructions for binding and elution. Eluted proteins were dialyzed against phosphate-buffered saline (PBS), followed by PBS plus 20% glycerol and stored at -20°C.

**Bottom-up proteomics**. LC-MS/MS proteomics was performed on an Orbitral Astral (Thermo) using Data Independent Acquisition, in biological triplicate. Thermo raw files were processed in DIA-NN,[3] or MAXQUANT with MaxDIA.[4] At least 2 peptides were required for protein identification. Data was searched against the reference proteome from *Corynebacterium diphtheriae* NCTC 13129 (UP000002198 Uniprot Proteome ID, access date December 4$^{th}$, 2025).[5] Gel-enhanced LC-MS/MS (GeLC-MS) proteomics fractions were acquired in DDA mode on a Q-exactive plus orbitrap mass spectrometer (Thermo) using data-dependent acquisition, 60 minute gradient.

**Protein purification for *C. diphtheriae* growth studies.** Recombinant proteins for *C. diphtheriae* feeding experiments were prepared as previously described.[6] In brief, ChtA$^{CR}$ (residues 112-291) or IsdA$^{NEAT}$ (residues 58-188) was expressed from the pET-28b-based plasmid containing an N-terminal small ubiquitin-like modifier (SUMO) fusion tag (pSUMO) and a 6x-His purification tag in E. coli BL21 (DE3) chemically competent cells (New England Biolabs, Beverly, MA). Cells were grown in LB media at 37 °C to a final OD600 of 0.6-0.8 before addition of 500 µL of 8 mM hemin chloride and induced with 1 mM isopropyl-β-D-thiogalactopyranoside (IPTG) followed by overnight growth at 17 °C. Cell pellets were harvested, resuspended, and lysed in lysis buffer (50 mM sodium phosphate, 300 mM NaCl, pH 7.0) supplemented with 1 mg/mL egg white lysozyme, 2 mM phenylmethylsulfonyl fluoride (PMSF, Sigma), a protease inhibitor cocktail (Sigma-Aldrich), and 0.5 mg Serratia marcescens nuclease.[7] His-tagged protein was purified from centrifuge-clarified lysate by passage over a HisPur Co2+ column (Thermo Fisher Scientific) followed by additional washes and elution with high imidazole buffer. The SUMO fusion was removed by adding the ULP1 protease (purified in-house) and second passage through a HisPur Co2+ column. Size-exclusion chromatography was used as a final purification step using an Akta FPLC system (GE Healthcare Life Sciences) equipped with a Superdex S75 size exclusion column. Hemophore proteins initially copurified with mixed amounts of endogenous

*E. coli* heme; fully holo proteins were obtained by incubation with 20-fold molar excess hemin chloride dissolved in 0.1 M NaOH with agitation for 16 hours. Separation of excess hemin was achieved through the use of DEAE-Sepharose anion exchange resin.[8]

***C. diphtheriae* growth studies.** C. *diphtheriae* (strain 1737) cells were grown in 96-well, 200 µL microtiter plate in mPGT media to limit iron access.[1] Overnight cultures were grown in 2 mL heart infusion broth (Difco) with 0.2% Tween 80 (Millipore Sigma) (HIBTW) with shaking before 1:2 dilution with fresh HIBTW the next morning. Cells were allowed to grow for an additional 1-2 hours, after which 500 µL of culture was harvested and spun down at 13,000 ×g for 1 minute. Cell pellets were resuspended in 1 mL mPTG supplemented with 1 µM FeCl3 before being allowed to grow for 4-6 hours at 37°C with shaking, in order to iron-stress the bacteria. These cultures were used to seed 200 µL wells of mPTG with 30 µM EDDHA, to a starting OD600 of 0.02. Blood-purified MetHb, recombinant holo-ChtA$^{CR}$, or recombinant holo-IsdA$^{NEAT}$ were supplemented 0.6 µM on heme-basis in triplicate. OD600 was measured for 24 hours at 37°C every 10 minutes using a SpectraMax iD3 plate reader (Molecular Devices).

#### Supplementary Figures

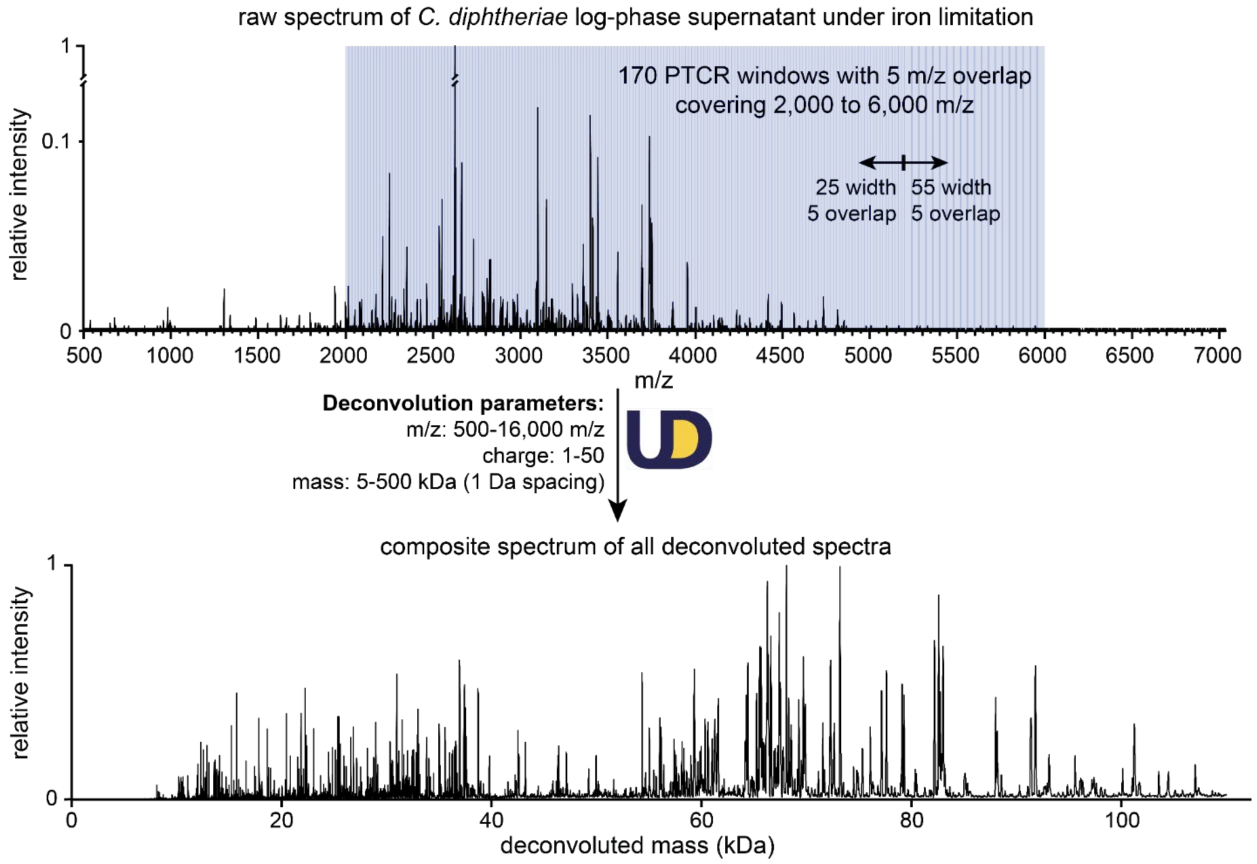


**Figure S1**: Data independent acquisition charge-reduction using proton transfer charge reduction (PTCR) of *C. diphtheriae* exoproteome collected on a Thermo Scientific Orbitrap Ascend BioPharma Tribrid mass spectrometer with Native MS option. (A) Raw mass-to-charge spectrum from *C. diphtheriae*'s iron-limited native exoproteome, indicating isolation windows used for PTCR. (B) Composite sum of all deconvolved normalized mass spectra. Individual PTCR spectra from isolation windows indicated in (A) were deconvolved in Unidec according to the Unidec deconvolution parameters indicated by the arrow and Unidec logo. This resulted in a total of 371 proteoforms with high-confidence (defined by having a Unidec D-score greater than 0.4 and normalized intensity greater than 3% in at least 2 of the 175 windows). Peak spacing was 23 Da to mitigate addition of ammonium and sodium proteoform adducts to the dataset.

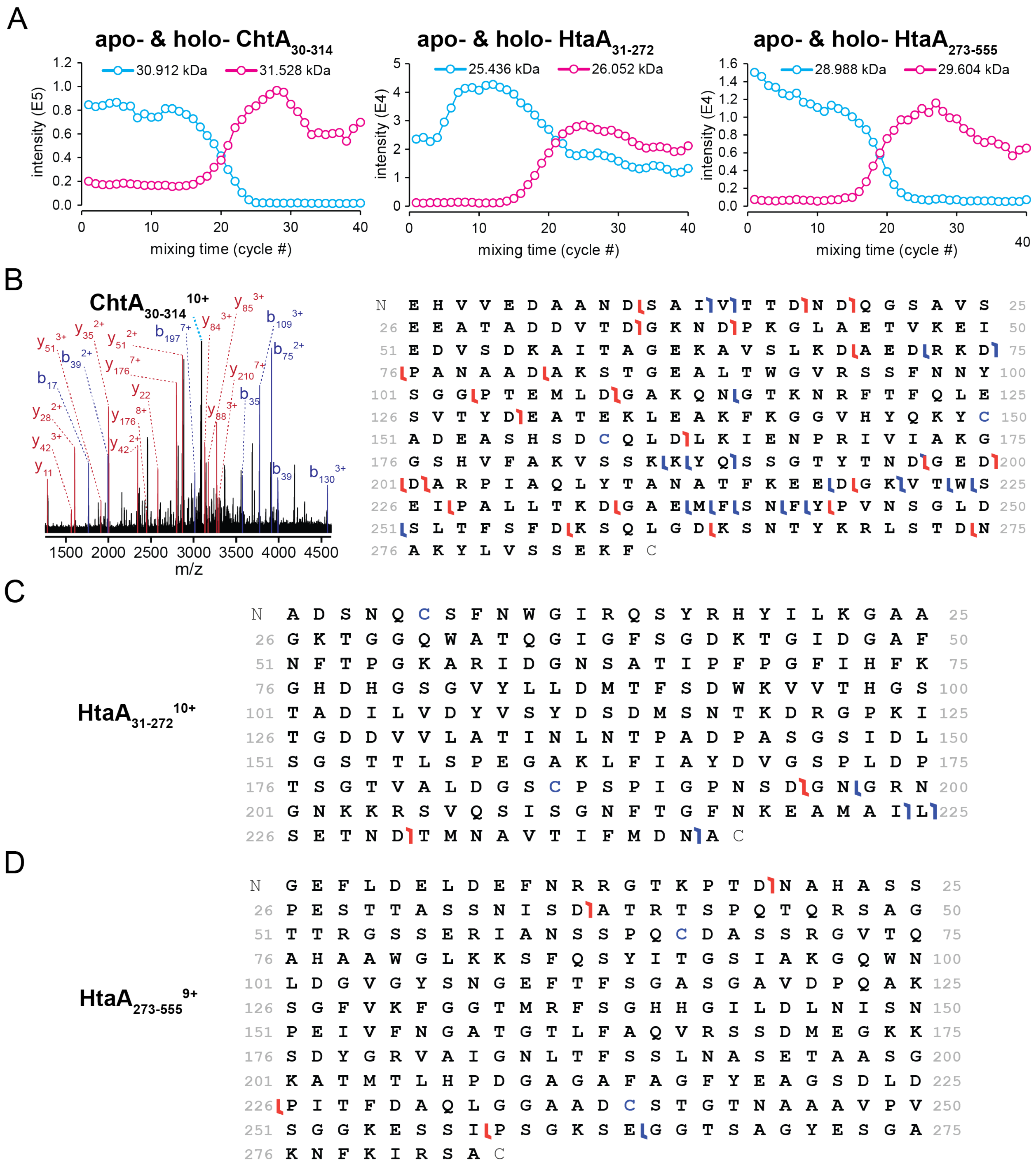


**Figure S2**: ProteoMIX verification of ChtA and HtaA hemophores ($ChtA_{30-314}$, $HtaA_{31-272}$, and $HtaA_{273-555}$). (A) Intensities across ProteoMIX mixing time for apo- and holo- heme-bound forms of $ChtA_{30-314}$, $HtaA_{31-272}$, and $HtaA_{273-555}$. Note that a reciprocal change in intensity is observed for the apo- and holo-forms of each hemophore, and their magnitude is similar. (B) Native top-down (nTD) higher-energy collisional dissociation (HCD) fragmentation spectra of the 10+ charge-state of apo-$ChtA_{30-314}$ (3092 m/z) at 110V (left). The cleavage map is shown on the right. (C) Cleavage map of the 9+ charge state of apo-$HtaA_{31-272}$ proteoform (2827 m/z) from nTD HCD fragmentation at 110 V.

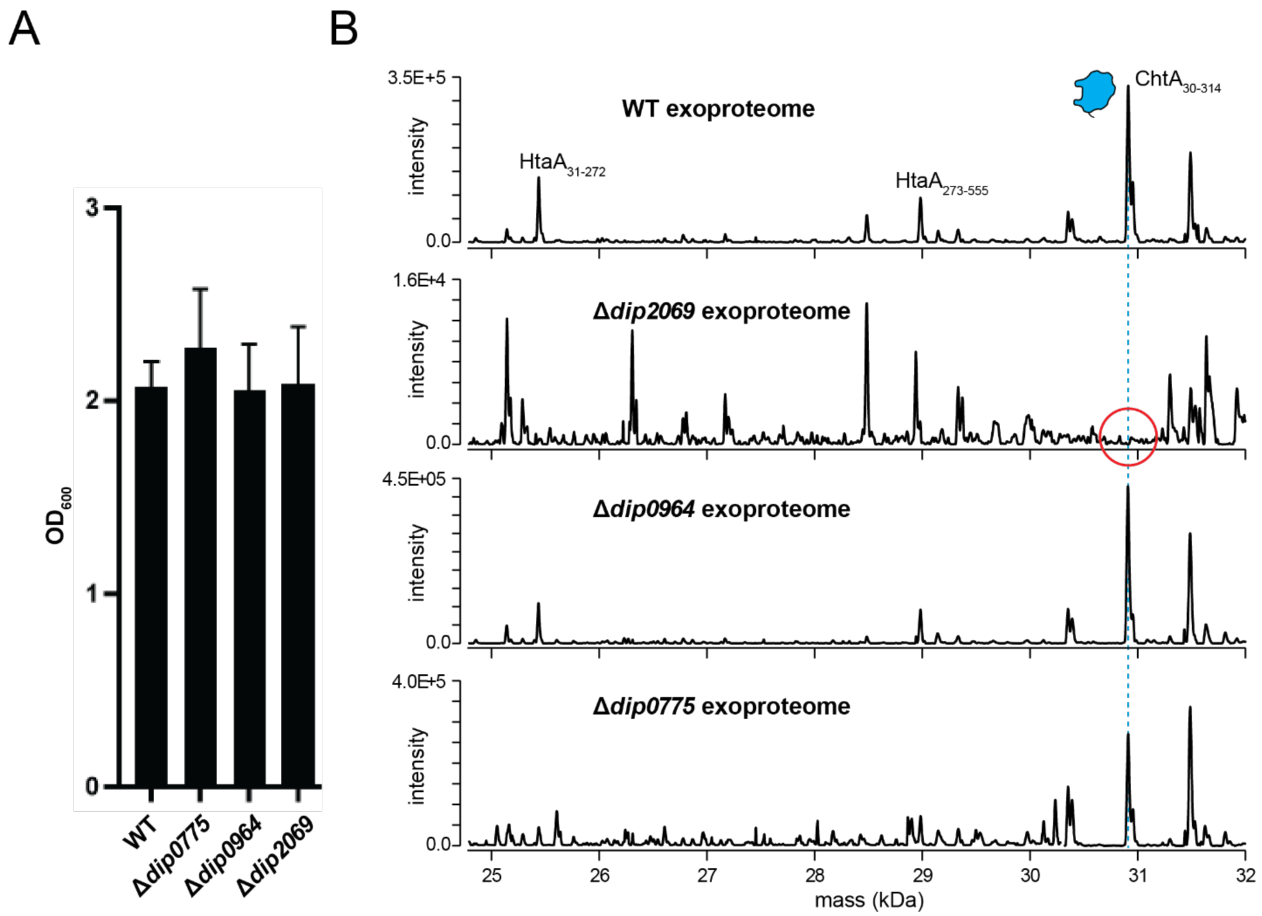


**Figure S3**: Growth and detection of secreted truncated heme-binding proteoforms in WT, *Δdip2069*, *Δdip0964*, and *Δdip0775 C. diphtheriae* mutants. (A) Terminal OD600 growth values of each strain in chemically defined mPGT media with 0.3 μM $FeCl_3$ supplementation. No significant difference is observed across strains. (B) Mass vs intensity spectrum of each strain from DIA-charge reduction of each proteome. Dotted lines track the mass of each hemophore identified from ProteoMIX in the WT exoproteome ($ChtA_{30-314}$, $HtaA_{31-272}$, and $HtaA_{273-555}$). All three hemophores are clearly observed in WT, *Δdip0964*, and *Δdip0775 C. diphtheriae* mutants. The Δ2069 exoproteome is considerably different from the others, with no peak corresponding to $ChtA_{30-314}$, and very low peak intensities observed at masses corresponding to $HtaA_{31-272}$, and $HtaA_{273-555}$.

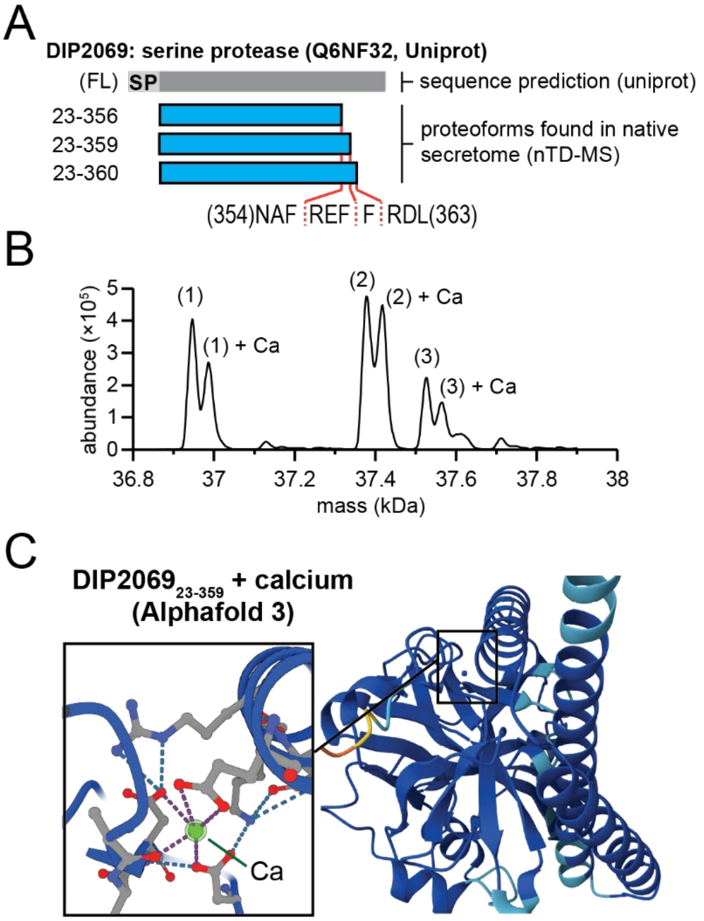


**Figure S4**: Calcium binding to native DIP2069 proteoforms. (A) Schematic of DIP2069 showing the proteoforms identified by nTD-MS. Figure shows DIP2069 predicted gene product from Uniprot Accession, Q6NF32. The identified proteoforms from nTD-MS are shown in blue. The N- and C-terminal amino acid numbers are shown on the left, and the C-terminal cleavage sites are shown on the bottom. Note that amino acids N-terminal to the signal peptide ("SP") signal peptidase I cleavage site are missing in these truncated proteoforms, consistent with secretion through the Sec translocon. (B) Each proteoform contains an adduct approximately 40 Da higher than the apo-form, consistent with calcium binding. Calcium is an established co-factor for serine proteases that adopt a chymotrypsin-like fold. (C) Alphafold 3 model of DIP2069$_{23\text{-}359}$ with calcium showing high confidence (>0.9) of the chymotrypsin-like fold.

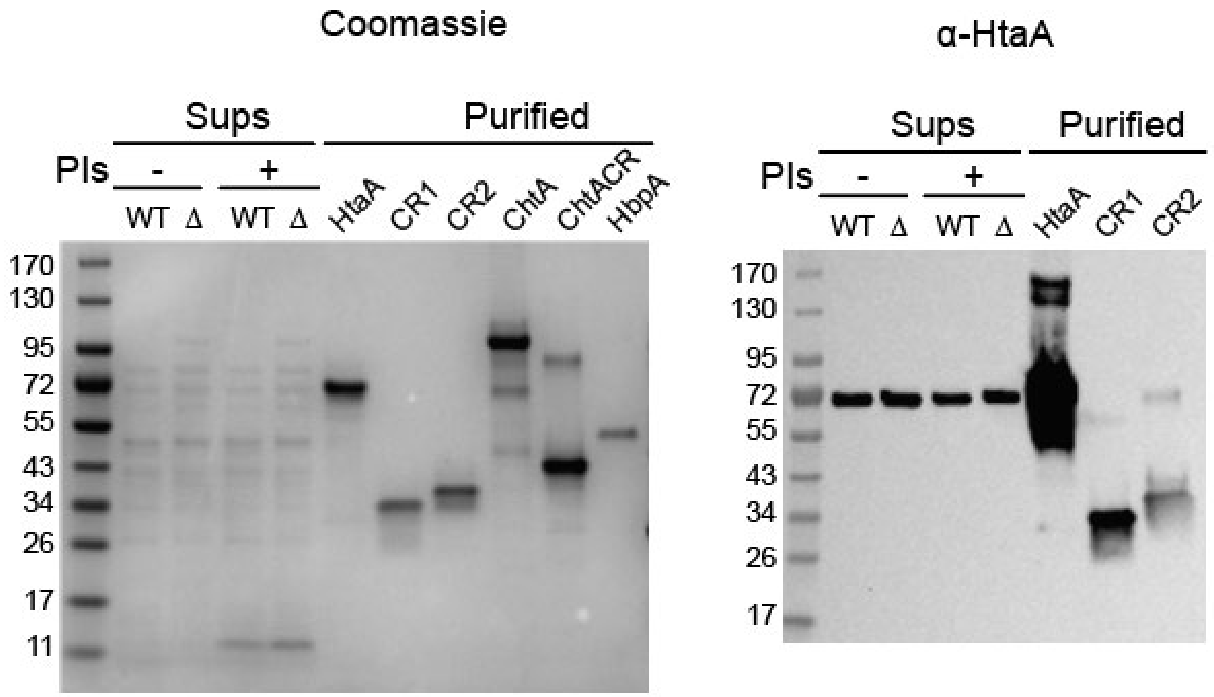


**Figure S5**: Western blot analysis on WT and Δ*dip2069* ("Δ") C. diphtheriae exoproteomes from stationary-phase iron-limited media cultures. A "+" indicates presence of protease-inhibitor cocktail addition to exoproteome immediately following cell pelleting, like native exoproteome preparation, and "-" indicates no addition whatsoever. Anti-HtaA antibodies show no change in HtaA-FL abundance in WT vs Δ*dip2069* exoproteomes.

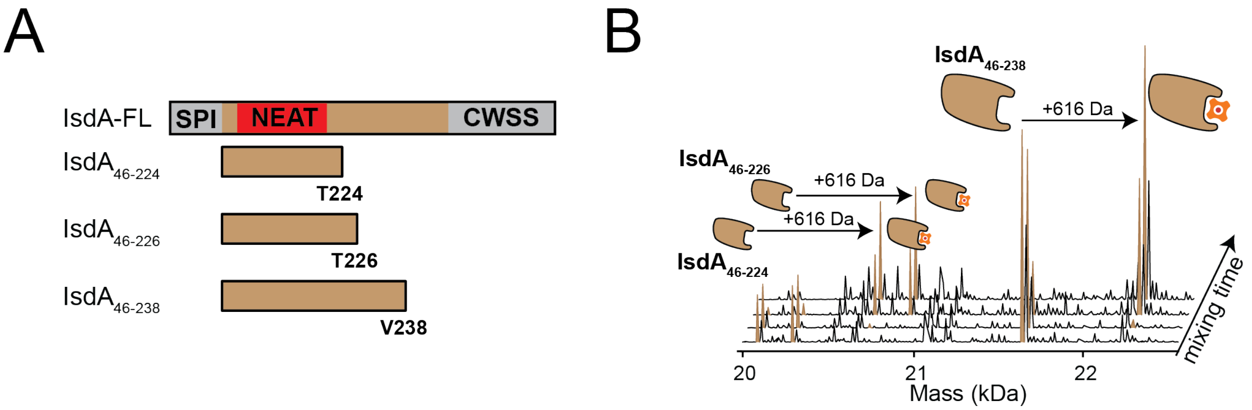


**Figure S6:** SLOMO-nMS detects heme binding to truncated *Staphylococcus aureus* IsdA NEAT domain-containing proteoforms. (A) Schematic of full-length IsdA (IsdA-FL) showing the signal peptide (SPI), NEAT domain, and C-terminal cell-wall sorting signal (CWSS) which contains an LPxTG motif, and the three soluble constructs analyzed by native MS ($IsdA_{46-224}$, $IsdA_{46-226}$, and $IsdA_{46-238}$; C-terminal residues indicated). (B) ProteoMIX deconvolved mass spectra showing heme binding to IsdA truncations in (A). Desalted *S. aureus* exoproteome from 16-18 hour stationary phase low-iron (RPMI 1640) cultures were concentrated, buffer-exchanged into 200 mM ammonium acetate, and subject to ProteoMIX workflow to find search for hemophores capable of extracting heme from Hb. Spectra are shown as deconvolved mass distributions; apo species are observed initially, followed by appearance of a +616 Da mass shift corresponding to formation of the 1:1 heme-bound complex (brown cartoons).

### Supplementary Tables

##### Table S1. Proteoforms identified in the exoproteome of *C. diphtheriae* using native top-down MS.

| Gene Name | Gene Name (Ordered Locus) | UniProt Accession Code | Protein Name | Proteoform Identified (nTD-MS) | % exoproteome by LC-MS/MS (bottom-up) |
|---|---|---|---|---|---|
| **DIP2069** | DIP2069 | Q6NF32 | Secreted protease | (SP)\|-359(F) | 19.74 ± 8.1% |
| **DIP2069** | DIP2069 | Q6NF32 | Secreted protease | (SP)\|-356(F) | 19.74 ± 8.1% |
| **DIP2069** | DIP2069 | Q6NF32 | Secreted protease | (SP)\|-360(F) | 19.74 ± 8.1% |
| **Tox** | DIP0222 | Q6NK15 | Diphtheria toxin | (SP)\|-end | 10.14 ± 1.9% |
| **Tox** | DIP0222 | Q6NK15 | Diphtheria toxin | (SP)\|-end + 980 Da | 10.14 ± 1.9% |
| **ChtA** | DIP1520 | Q6NGJ3 | Heme binding protein | (SP)\|30-314(F) | 7.55 ± 1.5% |
| **HtaA** | DIP0625 | Q6NIZ1 | Heme binding protein | (A)\|273-555(A) | 5.77 ± 1.2% |
| **HtaA** | DIP0625 | Q6NIZ1 | Heme binding protein | (SP)\|-272(A) | 5.77 ± 1.2% |
| **SlpA** | DIP0365 | Q6NJN3 | Trehalose corynomycolyl transferase | (SP)\|-end | 4.89 ± 0.7% |
| **FrgD** | DIP1062 | Q6NHR9 | Iron-siderophore uptake system exported solute-binding component | (F)\|55-end | 2.98 ± 0.4% |
| **FrgD** | DIP1062 | Q6NHR9 | Iron-siderophore uptake system exported solute-binding component | (R)\|54-end | 2.98 ± 0.4% |
| **MntA** | DIP0615 | Q6NJ01 | ABC transport system exported protein | (SPII)-end; +154 Da modification | 1.56 ± 0.1% |
| **RpfA** | DIP0775 | Q6NIK0 | Resuscitation-promoting factor Rpf1 | (SP)\|-190\|(F) | 1.18 ± 1.1% |
| **RpfA** | DIP0775 | Q6NIK0 | Resuscitation-promoting factor Rpf1 | (SP)\|-end | 1.18 ± 1.1% |
| **SprX** | DIP0964 | Q6NI15 | Protease | (SP)\|-238(A) | 0.86 ± 0.2% |
| **DIP0013** | DIP0013 | Q6NKK5 | Long Rib domain-containing protein | (SP)\|-end | 0.73 ± 0.1% |
| **DIP0350** | DIP0350 | Q6NJP6 | Secreted protease | (SP)\|-end | 0.7 ± 0.3% |
| **DIP1960** | DIP1960 | Q6NFD4 | Exported protein | (SP)\|-end | 0.68 ± 0.1% |
| **LpqC** | DIP1116 | Q6NHM0 | Exported esterase/hydrolase | SP 33-end344 | 0.48 ± 0% |
| **RplL** | DIP0437 | Q6NJG5 | 50S ribosomal protein L7/L12 | FL | 0.34 ± 0.1% |
| **Pgk** | DIP1309 | P62411 | Phosphoglycerate kinase | Nme-end | 0.32 ± 0.1% |
| **FadD1** | DIP2190 | Q6NES7 | Polyketide synthase subunit | FL | 0.27 ± 0.1% |
| **PpiA** | DIP0025 | Q6NKJ3 | Peptidyl-prolyl cis-trans isomerase | FL | 0.26 ± 0.1% |
| **PpiA** | DIP0025 | Q6NKJ3 | Peptidyl-prolyl cis-trans isomerase | Nme-end | 0.26 ± 0.1% |
| **DIP0878** | DIP0878 | Q6NI99 | ABM domain-containing protein | FL (dimer) | 0.24 ± 0.1% |
| **TrxA** | DIP2284 | Q6NEJ0 | Thioredoxin | Nme-end | 0.23 ± 0.04% |
| **Tsf** | DIP1507 | P61332 | Elongation factor Ts | FL | 0.19 ± 0.04% |
| **Tsf** | DIP1507 | P61332 | Elongation factor Ts | Nme-end | 0.19 ± 0.04% |
| **TpiA** | DIP1308 | Q6NH37 | Triosephosphate isomerase | Nme-end | 0.17 ± 0.01% |
| **TpiA** | DIP1308 | Q6NH37 | Triosephosphate isomerase | Nme-end (dimer) | 0.17 ± 0.01% |

| | | | | | |
|---|---|---|---|---|---|
| **AnsA** | DIP0491 | Q6NJB8 | Secreted amino acid hydrolase | (SP)\|-end (dimer) | 0.17 ± 0.04% |
| **Ndk** | DIP1783 | Q6NFV3 | Nucleoside diphosphate kinase | Nme-end (hexamer) | 0.17 ± 0.02% |
| **DIP2017** | DIP2017 | Q6NF77 | Secreted protein | [31-423] (sparse N-term) | 0.16 ± 0.03% |
| **AhpD** | DIP1419 | Q6NGT4 | Alkyl hydroperoxide reductase AhpD | Nme-end (trimer) | 0.14 ± 0.02% |
| **DirA** | DIP1420 | Q6NGT3 | Alkyl hydroperoxide reductase C | Nme-end | 0.12 ± 0.04% |
| **PckG** | DIP2180 | Q6NET5 | Phosphoenolpyruvate carboxykinase [GTP] | Nme-end | 0.1 ± 0.02% |
| **DIP0571** | DIP0571 | Q6NJ42 | Secreted protein | (SP)\|-end | 0.08 ± 0.04% |
| **PepO** | DIP0154 | Q6NK83 | Endopeptidase | (SP)\|-end | 0.074 ± 0.004% |
| **Frr** | DIP1505 | P61303 | Ribosome-recycling factor | FL | 0.07 ± 0.001% |
| **CtaC** | DIP1629 | Q6NG98 | Cytochrome c oxidase subunit 2 | (A)\|138-end | 0.07 ± 0.03% |
| **DIP0793** | DIP0793 | Q6NII2 | Secreted protein | (SP)\|-end | 0.05 ± 0.03% |
| **Ppa** | DIP2006 | Q6NF88 | Inorganic pyrophosphatase | Nme-end (hexamer) | 0.04 ± 0.01% |
| **OdhI** | DIP1204 | Q6NHD4 | Oxoglutarate dehydrogenase inhibitor | Nme-end | 0.04 ± 0.01% |
| **Tal** | DIP1303 | Q6NH42 | Transaldolase | Nme-end | 0.03 ± 0.01% |
| **FkbP** | DIP0786 | Q6NII9 | Peptidyl-prolyl cis-trans isomerase | FL | 0.02 ± 0.01% |
| **Mce** | DIP1057 | Q6NHS4 | Methylmalonyl-CoA epimerase | Nme-end | 0.016 ± 0.003% |
| **Mce** | DIP1057 | Q6NHS4 | Methylmalonyl-CoA epimerase | Nme-end (dimer) | 0.016 ± 0.003% |
| **Efp** | DIP1340 | Q6NH07 | Elongation factor P | Nme-end | 0.01 ± 0.01% |
| **ProS** | DIP1482 | Q6NGM7 | Proline--tRNA ligase | FL (dimer) | 0.01 ± 0.002% |
| **DIP0711** | DIP0711 | Q6NIQ6 | DUF4288 domain-containing protein | NAc-end (dimer) | 0.01 ± 0.01% |
| **DIP0761** | DIP0761 | Q6NIK8 | Cupin type-2 domain-containing protein | Nme-end (dimer) | <0.01 |
| **DIP0701** | DIP0701 | Q6NIR6 | HAD family hydrolase | FL | <0.01 |
| **Mdh** | DIP1787 | P61974 | Malate dehydrogenase | Nme-end | <0.01 |
| **DIP0615** | DIP0615 | Q6NJ01 | ABC transport system exported protein | X-end | <0.01 |

The proteoform gene names, gene names corresponding to their ordered locus, Uniprot accession numbers, protein names, identified proteoform, and % iBAQ abundance in the exoproteome determined using LC-MS/MS bottom-up proteomics. Proteoform annotations starting with “(SP)” or “(SPII)” indicate an N-terminus in agreement with the signal peptide peptidase I or II cleavage, respectively, determined from SignalP 6.0.[9] Numbers indicate novel truncations, and parentheses next to the number indicate the P1 residue giving rise to the associated truncation. A vertical bar, “|”, is used to illustrate the cleavage site between the identified proteoform and its N-terminal processed residue(s) such as signal peptide cleavage sites or novel cleavage sites indicated by a P1 residue. N-terminal methionine cleavages are indicated by “Nme”. An “X” indicates that the terminus of the protein is unidentified. And other post-translations modifications are described after the semicolon.
